# Optimizing irrigation during heat events sustains grapevine physiology and fruit production

**DOI:** 10.64898/2026.02.10.705159

**Authors:** Martina Galeano, Andrew J. McElrone, Lauren E. Parker, Nicolas Bambach, Luis Sanchez, Nick Dokoozlian, Sophia Bagshaw, Paul Bringas, Kayla Elmendorf, Elisabeth J. Forrestel

## Abstract

1. Increasing frequency, intensity, and duration of heat waves (HWs) threaten agricultural production globally by constraining physiological function and fruit production. Supplemental irrigation mitigates heat stress in grapevine and other woody perennial crops, yet water scarcity necessitates optimized irrigation strategies during extreme heat.
2. We conducted a three-year field trial in a commercial Cabernet Sauvignon vineyard, applying differential irrigation only before and during naturally occurring HWs: baseline (50% ET), moderate (90-120% ET), and high (120-180% ET). We monitored water potentials, leaf gas exchange, canopy temperature, yield, and berry composition.
3. Baseline irrigation consistently reduced net photosynthesis, stomatal conductance, and leaf cooling capacity during HWs. Moderate supplemental irrigation maintained gas exchange, transpiration, and leaf temperature, mitigating yield losses. Excessive irrigation beyond moderate levels provided no additional physiological benefit and decreased crop water use efficiency and berry quality.
4. Our results demonstrate that targeted, event-based irrigation sustains grapevine physiological performance and fruit production under extreme heat, whereas both insufficient and excessive water negatively affect carbon assimilation, stomatal regulation, and crop productivity. These findings emphasize the importance of aligning water management with heat event timing to preserve vine function, optimize water use, and maintain yield and fruit quality in water-limited regions.

## Introduction

Long-lived perennial woody crops are increasingly exposed to extreme heat waves (HWs) of higher intensity, frequency and duration (Diffenbaugh *et al*., 2021; Yin *et al*., 2022; Lee *et al*., 2023). These extreme weather events, typically defined as three or more days with anomalously maximum temperatures relative to a historical average for a given time of year (Robinson, 2001) can be particularly damaging to woody perennial crops, such as grapevines. High temperatures can arrest crop development, deplete carbohydrate reserves, decrease yield and quality, and ultimately induce early senescence of perennial crops (Sadras & Moran, 2013; Parker *et al*., 2020). Yet we have limited knowledge of how these crops will be impacted as most studies have been focused on annual crop responses (Parker *et al*., 2020; Breshears *et al*., 2021). While grapevines are known for tolerating low water availability and warm temperatures, it is critical to understand how the timing and amount of water application during HWs impact vine physiology and productivity as heat stress events become more frequent (Breshears *et al*., 2021; Coupel-Ledru *et al*., 2024) and are often coupled with soil and atmospheric drought.

Grapevine photosynthesis is stimulated and favored with rising temperatures until 25 to 30° C (although this varies by cultivar) (Kriedemann, 1968), but beyond this temperature a decrease in the photosynthetic performance, generally associated with stomatal limitation, is observed (Gouot *et al*., 2019b; Greer & Weston, 2010; Greer & Weedon, 2013). However, exposure to an extended period of extreme heat (e.g., a 14 day heat wave in Australia in 2009), causes a decrease of photosynthesis, mainly related to biochemical constraints such as carboxylation and ribulose-1,5-biphosphate limitation (Greer & Weedon, 2012, 2013). Exposure to extreme heat from fruit set through ripening has also been found to reduce berry size and total yield (Soar *et al*., 2009; Greer & Weston, 2010); (Coombe & Iland, 2004). Despite previous work in both natural and agricultural systems, there is a lack of understanding of how the timing of water availability mitigates impacts of extreme heat events on plant or crop performance and productivity.

Extreme temperatures generally result in higher water loss and resulting stress on the plant. Thus, irrigation can be used to mitigate the adverse effects of HWs by modifying vine water status and increasing evaporative cooling. Vines depend on transpiration (E) and evaporation from the ground to cool the leaves (Kool *et al*., 2014), and some cultivars have been shown to increase stomatal conductance (g_s_) under heat, allowing a higher transpiration rate under well watered conditions (Soar *et al*., 2009; Sadras *et al*., 2012). While there is evidence that warming has increased quality in some regions, most wine growing regions are currently at the optimal climatic conditions for the grape varieties cultivated there (Jones *et al*., 2005). Therefore, rising temperatures will likely alter the physiological performance of different grapevine varieties, potentially affecting growth, metabolism, and reproductive output. (Morales-Castilla *et al*., 2020; Sgubin *et al*., 2023; Coupel-Ledru *et al*., 2024; Parker *et al*., 2024).

Warm, dry summers and cold, wet winters are typical of Mediterranean climates, which are widely used for the cultivation of perennial woody crops such as grapevines (Gershunov & Guirguis, 2012). The average annual global temperature has increased dramatically (Bedsworth *et al*., 2018), but previous research suggests that changes in mean temperatures would have a minor effect on grape growing conditions (Jones & Davis, 2000). Yet, the associated increase in extreme temperatures like those that occur during HWs, coupled with increases in evaporative demand, represents a major constraint on the physiological performance and production of perennial crops under field conditions (White *et al*., 2006; Gershunov *et al*., 2010; Grossiord *et al*., 2020; van Leeuwen *et al*., 2024). Modeled end of century predictions in California of the frequency of three consecutive days with maximum temperatures above 38° C project a severe increase in the number of HWs (Figure 1 A & B). Rising temperatures will also shift winter precipitation patterns from snow to rain. This, combined with the majority of California’s annual precipitation taking place during winter, will increase the gap between the time when water is supplied by rivers and when grapevines start demanding water (Keller, 2010; Parker *et al*., 2020), which has major implications for agricultural areas that depend on snowmelt runoff (Qin *et al*., 2020). Furthermore, vines will be exposed to higher water stress due to an increase in evapotranspiration and water vapor pressure deficit (VPD) caused by exposure to higher temperatures and greater atmospheric water demand (Manabe *et al*., 2004; Grossiord *et al*., 2020; Albano *et al*., 2022; Fang *et al*., 2022). Collectively, these changes will increase water stress during the grapevine growing season, placing greater demand on increasingly scarce water resources to maintain physiological function (Keller, 2010; Parker *et al*., 2020).

**Figure 1.**
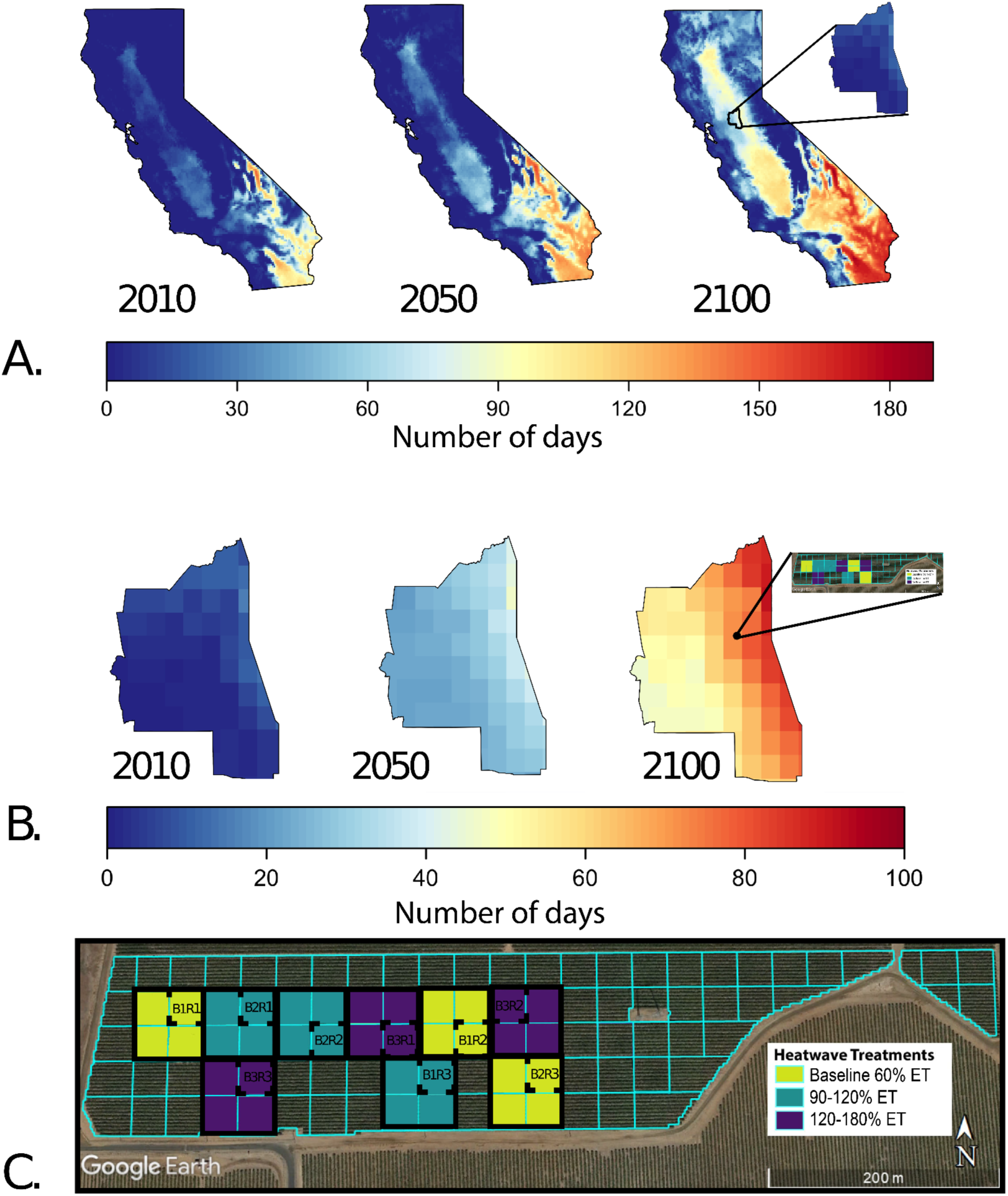
Changes in the number of days above the 38 degree Celsius threshold to be considered an extreme heat day for A. California, and B. the Lodi AVA, by mid-century (2050) and end of the century (2100) projections relative to a 1980-2010 normal, based on a model ensemble from the RCP 8.5 emissions scenario (Abatzoglou & Brown, 2012). C. Layout of the vineyard study site and experimental design of irrigation treatments, with the 30 x 30 meter squared 2019 subplots indicated with a dashed line within the 60 x 60 meter squared 2020 and 2021 subplots.

The objective of this study was to evaluate the impact that different amounts of irrigation carried out prior to and during HWs in a field experiment have on vine physiology, yield, and berry composition. Results from this study facilitate a better understanding of how water availability during HWs modulates physiological responses to extreme temperatures, and impacts water and crop use efficiency. Further, understanding how heat events and water availability ultimately influence the production of primary metabolites in fruit will provide insights into plant functional responses under thermal stress.

## Materials & Methods

### Experimental Design

This study was carried out during the 2019, 2020 and 2021 growing seasons in a commercial vineyard located in Lodi, CA, one the most productive grape growing regions of the Central Valley (Figure 1C). The site is a 23 acre block of 9-year-old *Vitis vinifera* cv. Cabernet Sauvignon grafted onto 1103 Paulsen with a high wire bilateral cordon trellis system. Rows are oriented East-West with a vine spacing of 1.8 m and a row spacing of 3 m.

The vineyard block was divided into 98 subplots of 30 m x 30 m with a variable rate drip irrigation (VRDI) system allowing the application of a differential irrigation to each individual subplot (Sanchez *et al*., 2017). Applied irrigation was scheduled throughout the growing season using a method that calculates crop evapotranspiration (ET_c_) using a combination of both ET_o_ from a nearby weather station and a dynamic, modified crop coefficient (K_c_) approach (Allen *et al*., 1998). K_c_ values were derived by using remotely-sensed normalized difference vegetation index (NDVI) specific to each 30 x 30 m variable rate zone, along with empirical parameters representing the expected water demands for a non-stressed vineyard.

### Defining Heatwaves & Irrigation Treatments

To assess current grower practices in response to predicted HWs, we conducted a survey of 58 growers across California. (<u>link to survey results</u>; Table S1 & <u>Summary of results</u>). The climate or weather-related issues that were reported to be the most concerning (“moderate”, “serious”, or “devastating”) were drought (48/58 respondents) and extreme daytime temperatures (46/58 respondents). Further, while not all respondents felt increasing irrigation was always an effective strategy, over 80% of respondents would elect to increase irrigation before and/or during a heat event with temperatures reaching over 100 ° F/38 ° C. Our experimental design for the study was thus directly informed by current commercial vineyard management practices in response to heat extremes.

In the context of this experiment, a HW was identified based on historical weather data and trends as a minimum of three consecutive days with daily maximum temperatures above 38° C. Recognizing that heat extreme parameters are defined relative to local historical conditions, the maximum temperature cutoff of 38° C was determined using the 1980-2010 climate as the baseline, the 95 % percentile of maximum temperatures during the growing season (April - October) were used to determine temperature above which was considered an extreme heat day. Maximum temperatures were recorded using a vineyard weather station located .8 kilometers from our trial at an adjacent experimental site (Kustas *et al*., 2022).

Differential irrigation treatments were applied only when a heat event took place and started one or two days before each HW and continued until the last day of the heat event. Three irrigation treatments were implemented during the 2019 and 2020 seasons: a control or baseline, which was exposed to deficit irrigation and held at 60% ET, a second treatment where the irrigation was double the baseline (120% ET) and third treatment with triple the amount of water of the baseline (180% ET). In 2021, irrigation treatments were adjusted to 60% ET, 90% ET, and 120% ET during HWs based on the growers’ realization, based on our findings, that excessive water was being used in the 3x treatment.

In 2019, three subplots (30 m x 30 m) per treatment were randomly selected and distributed along the site using the VRDI system to provide differential irrigation to each of them. In 2020 and 2021, each replicate was expanded to four subplots of 60 x 60 meter (Fig. 1C). Micrometeorological stations were deployed in each of the nine blocks to measure localized net radiation, air temperature, relative humidity, and canopy radiometric surface temperature. Soil sensors measuring volumetric water content, temperature, and electrical conductivity were inserted under the vines/drip irrigation emitters next to each of the stations at depths of 5, 30, 60, and 90-cm. Additional air temperature and relative humidity sensors were deployed (in one station per treatment) within the vine canopy with recordings taken at 15-minute intervals.

### Physiological measurements

Six randomly selected vines per subplot were selected for physiological measurements (N=18/treatment; Fig. 1C). Vines from the first and last two rows of each subplot, and the first and last two vines of each row were not sampled to prevent influence from surrounding subplots. Diurnal leaf-level gas exchange measurements and water potential measurements were taken bi-weekly from (pre-veraison) early July to the through harvest in 2019 and from flowering to mid September in 2020, as well as out prior to, during, and post-HW events. Only midday measurements were taken from flowering to mid September in 2021.

Three fully expanded sun-exposed leaves at a consistent developmental stage were monitored on each of the three vines per subplot (N=27 leaves per treatment). Gas exchange was measured using a Li-Cor 6800 portable photosynthesis system (Li-Cor, Lincoln, Nebraska, USA). Photosynthetic active radiation was set for each round of measurements during the day based on values given by an external quantum sensor (LI-190R). Air flow was set at 500 µmol m^-2^ s^-1^ and reference CO_2_ concentration was established at 400 ppm. Water potentials were taken using a Scholander pressure chamber (PMS Instrument Company, Albany, OR, USA). Ψ_leaf_ measurements were performed on the same leaves as gas exchange measurements during diurnals. For monitoring leaf temperature, recordings from the thermocouple located in the Li-Cor 6800’s leaf chamber were used, as well as Infrared Radiometer (IRT) MI-230 sensors (Apogee Instruments Inc., Logan, UT, USA).

### Berry sampling and juice chemical analysis

Berries were sampled bi-weekly from 50% veraison to harvest in 2019, and from fruit set to harvest in 2020 and 2021. Additionally, when a HW occurred, berries were sampled prior to, during, and post-heat event. Nine vines per subplot were randomly selected for sampling berries, following the same restrictions on vine selection as vines used for physiological measurements. Forty berries per vine (twenty berries on each side of the canopy) were collected from different clusters and sides of the cluster sampling a total of 360 berries per subplot treated for each sampling point along the ripening period.

Directly after sampling, a subset of 180 berries was divided into three replicates of sixty berries. Berries of each replicate were crushed, and the grape juice obtained was centrifuged at 4200 rpm for five minutes. Next, each juice sample was analyzed for TSS using a refractometer Sper Scientific 30051 (Sper Scientific LTD, Scottsdale, AZ, USA), pH with an Orion Star A211 pH meter (Thermo Fisher Scientific Inc., Waltham, MA, USA), and titratable acidity (TA) by titration with 0.1 N NaOH (VWR International, Radnor, PA, USA) with an end point at pH 8.2 (Iland *et al*., 2004).

### Yield and harvest

Grapes were considered to have reached maturity when total soluble solids (TSS) values were 25 ± 1° Brix. For 2019, six vines per subplot were hand-picked and harvested to estimate yield. Total number of bunches and total fruit fresh weight per vine was recorded. A total of three subplots per treatment were harvested. For 2020 and 2021, twenty-seven vines per subplot were harvested with a total of three subplots per treatment.

### Statistical analysis

All statistical analyses were carried out in R v 4.3.3 using tidyverse, ggplot2, ggpubr, cowplot, lubridate, dplyr, plyr, gtools, grDevices, agricolae, factoextra, ggpmisc, and magrittr packages. Code for all analyses is available at https://github.com/mgaleano-ucd/Borden_hills_heatwaves_complete.

Significant differences among treatments were tested by performing one-way analysis of variance for each sampling time and day, with p ≤ 0.05 (*), p ≤ 0.01 (**), and p ≤ 0.001 (***) as significance levels. Then, a post-hoc Tukey’s HSD test was carried out for mean comparison with p ≤ 0.05 used for minimum significance. Error bars and variability shown in tables and graphs are representative of the mean’s standard error (SE).

## Results

### Irrigation and plant water status

A total of nine heatwaves occurred at the study site across the three growing seasons, with two occurring in 2019, four in 2020, and 3 in 2021 (Fig. 2). The heatwaves were spread throughout the growing seasons occurring in May, June, July, August and September, which represented all stages of fruit development from the end of flowering through harvest. Differential irrigation treatments were applied only during the heatwaves (see gray bars in Fig. 2), and resulted in higher cumulative water application in the supplemental treatments compared to the baseline in all years (Fig. S1). Cumulative irrigation increased from 2019 to 2021 across all treatments with differences reflecting varied winter precipitation in each growing season (Fig. S1). For example, winter precipitation was significantly greater in 2019 and resulted in less overall applied water (Fig. S1), and a delayed start to irrigation due to soil water reserves (Fig. 2D). Conversely, the low rainfall in 2021 was connected with greater applied water across all treatments (Fig. S1).

**Figure 2.**
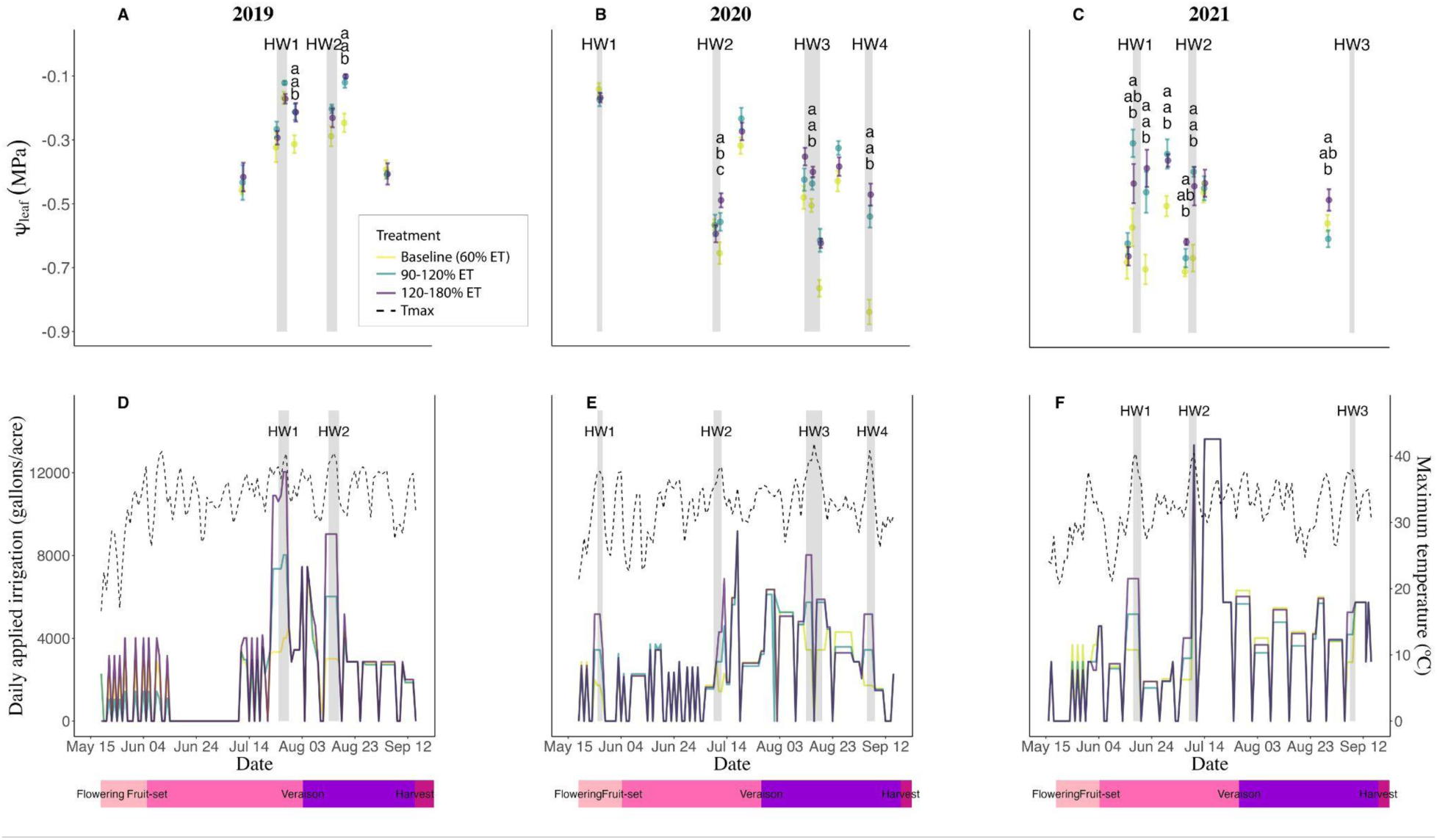
Predawn water potentials (Ψ _leaf_) from vines with three differential irrigation treatments during HWs, Baseline (60% ET), 90-120% ET, and 120-180% ET, collected during the 2019 (A), the 2020 season (B), and the 2021 season (C). Daily maximum temperature and total irrigation applied for different treatments in the 2019 (D), 2020 (E), and 2021 (F) growing seasons. Heatwave events (3 or more days predicted to have a maximum temperature above the 38 degrees Celsius threshold) are highlighted in gray. Error bars are SEM (n=9). P < 0.05; significance was evaluated by one-way analysis of variance and Tukey HSD was used to study post hoc differences among treatments, letters denote significant differences.

Differences imposed by the irrigation treatments were reflected in physiological changes in the vines. In terms of water relations, vines from each of the treatments exhibited similar predawn Ψ_leaf_ prior to the onset of differential irrigation during heatwaves. Predawn Ψ_leaf_ became generally lower (i.e. more negative) during and immediately following heatwaves in the baseline treatment compared to the supplement irrigation treatments, which is consistent with greater water stress (Fig. 2; Fig. S2). This pattern is very clear for the 3rd and 4th heatwaves in 2020, where predawn Ψ_leaf_ of the baseline was much lower than the other treatments. Overall, predawn Ψ_leaf_ values were higher (i.e., less negative) in 2019 compared to the other years consistent with high soil moisture reserves to start the season associated with greater winter precipitation. While the supplemental irrigation treatments were dialed back in 2021 to 1.5 and 2X the baseline, predawn Ψ_leaf_ was still significantly lower in the baseline compared to the supplemental treatments. In general, midday Ψ_leaf_ water potentials were less responsive to treatments (i.e., fewer significant differences) than midday Ψ_stem_ (Fig. S2. D-F). Midday Ψ_stem_ followed similar trends, albeit more negative values, as the predawn values, further corroborating the presence of greater stress in the baseline treatment (Fig. 2).

Midday Ψ_leaf_ followed a similar pattern with generally lower values in the baseline treatment when they did occur, but the differences between the treatments typically disappeared during the heatwave events, which is consistent with higher leaf transpiration in the supplemental treatments resulting in Ψ_leaf_ dropping to similar levels as the baseline at midday (Fig. S2). This pattern is particularly clear for midday Ψ_leaf_ in 2020, where the baseline treatment was consistently lower on average, but usually not significantly different than the other treatments (Fig. S2). During the 2020 and 2021 growing seasons, the changes to more negative water potential values from predawn to midday values were more pronounced in the supplemental irrigation treatments, due to increased rates of transpiration, stomatal conductance and latent cooling, with more plant available water. Yet, there was no additional benefit of applying excess water (beyond 1.5x baseline) in terms of water status stress reduction.

### Leaf-level gas exchange, temperature, and water use efficiency

Overall, A and g_s_ were significantly impaired in the baseline treatment relative to the 120% ET treatment, while there were no significant differences observed between the 120% and higher (180%). Significant decreases in net photosynthesis (A) and stomatal conductance (g_s_) were observed in the baseline treatment relative to the supplemental irrigation treatments (Fig. 3A, D) during the two heatwave events in 2019. For both A and g_s_, these differences persisted throughout heat waves and up until harvest, yet only differences in g_s_ were maintained through harvest. During 2020, the same significant trends were observed, but there was recovery between heat events earlier in the season (Fig. 3B, E) Overall treatment effects were less pronounced in 2021, but followed the same trends (Fig. 3C, F), likely due to the reduced difference between treatments in water application in 2021, and the overall higher amounts of irrigation applied during the season leading to less difference in water applied across treatments (Fig. S1). Leaf temperature in 2019 had a consistent response to stomatal closure seen in the baseline treatment when differential irrigation was implemented and was significantly higher in the baseline treatment compared to the supplementally irrigated treatments during both heat wave events (Fig. 3G). In 2020 and 2021, leaf temperature responses were less pronounced than 2019 (Fig. 3 H,I). During the 2020 season, the baseline treatment showed a significantly higher temperature only during HW4_2020_. In 2021, a treatment effect was only observed in two instances throughout the season; one prior to HW2_2021_, with a significantly higher temperature in the baseline compared to the other treatments, and one during HW3_2021_, with higher temperature between the baseline and the 90% ET treatment (1.5x baseline).

**Figure 3.**
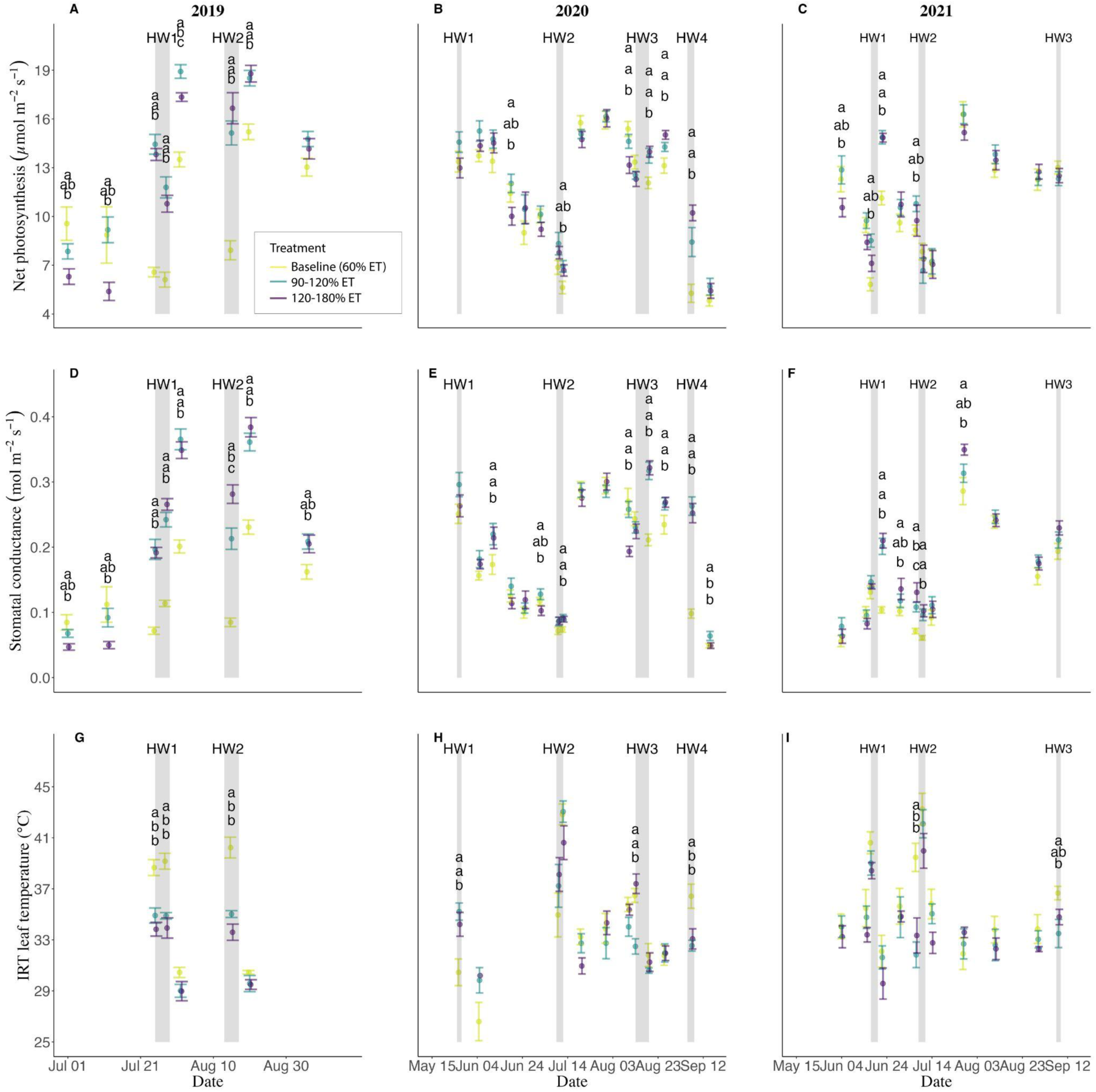
Midday measurements of net photosynthesis, A (top), stomatal conductance, g_s_ (middle), and IRT leaf temperature (bottom) carried out throughout the 2019 (A, D, G), 2020 (B, E, H), and 2021 seasons (C, F, I). Heatwave events (3 or more days predicted to have a maximum temperature above the 38 degrees Celsius threshold) are highlighted in gray. Error bars are SEM (n=27) for A and g_s_ and (n=9) for IRT leaf temperature. P < 0.05; significance was evaluated by one-way analysis of variance and Tukey HSD was used to study post hoc differences among treatments, letters denote significant differences.

Leaf-level intrinsic water use efficiency (iWUE, or A/g_s_) was significantly higher in the baseline treatment in 2019 during the first HW and through the remainder of the season. During the 2020 and 2021 growing seasons, there was the same trend of higher iWUE in the baseline treatment, but significance was constrained to HWs, and did not persist between events (Fig. 4B,C). Despite the trend of having higher intrinsic WUE at the leaf level all three years in the baseline treatment, that was not the case when looking at crop water productivity, defined as the ratio between yield and total irrigation applied. In the case of crop WUE, no significant differences between treatments were seen across the three years of the study, however, the 120% (in 2019 and 2020) and 90% ET (in 2021) were the treatment<u>s</u> that had on average the highest crop WUE each year (Fig. S3). Further, crop WUE was substantially higher in 2019 relative to 2020 and 2021, which follows the higher rates of net photosynthesis and stomatal conductance reached in 2019 in conjunction with greater available water in the soil (i.e., greater pre-dawn water potentials). Overall, heatwaves were less impactful in 2020 and 2021, likely as a result of substantial increases in irrigation across the vineyard and more similar amounts of seasonal irrigation applied (Fig. S1). Despite having more positive pre-dawn water potentials, the baseline treatment had more dramatic reductions in both net assimilation rates and stomatal conductance, reflected both in midday measurements and integration across the day (diurnal measurements; Fig. 4). For all years, supplemental irrigation beyond the 90-120% ET treatments provided no additional benefit in terms of gas exchange and leaf temperature responses.

**Figure 4.**
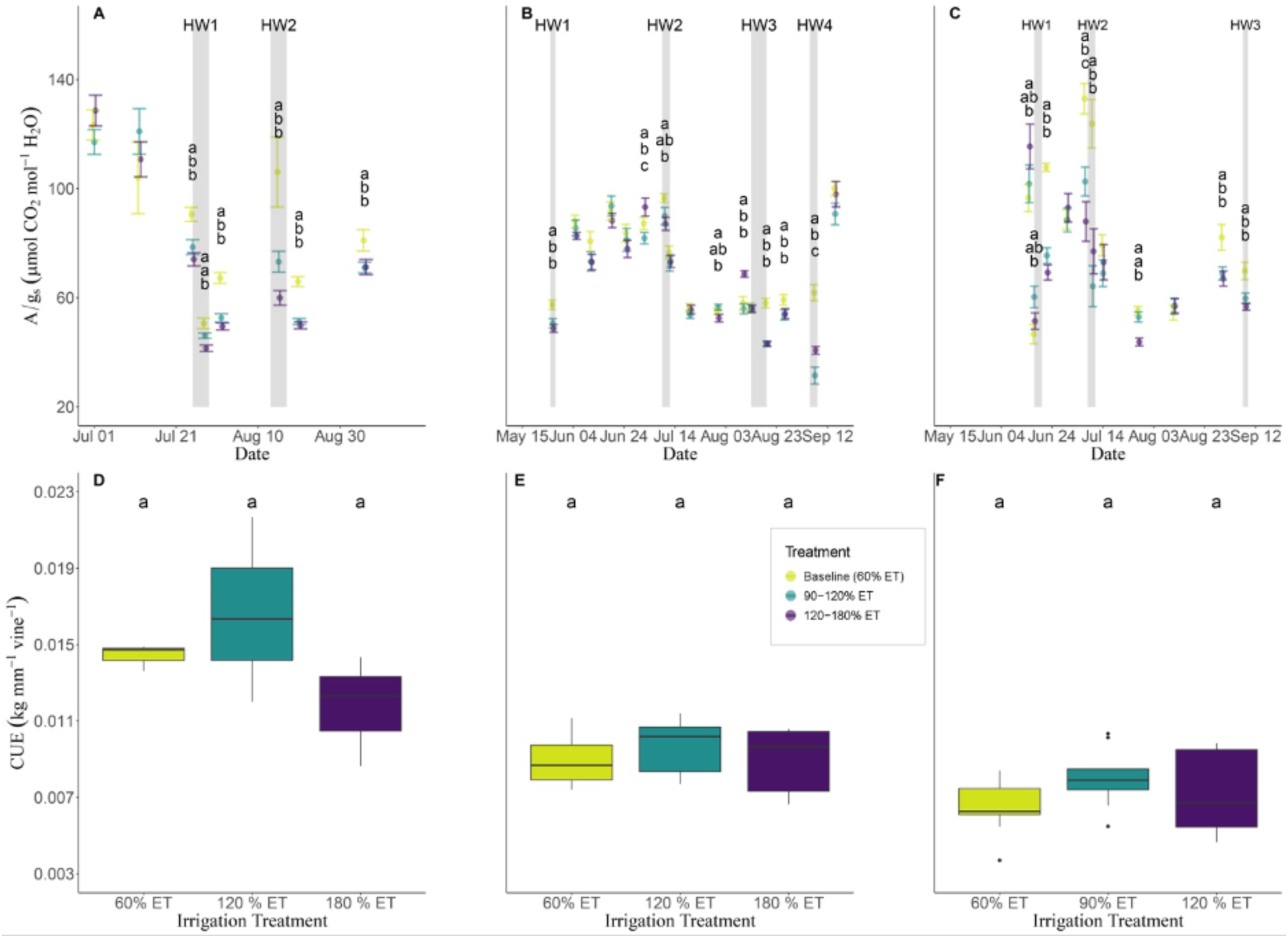
Midday measurements of intrinsic WUE, A/g_s_ carried out throughout the 2019 (A), 2020 (B), and 2021 seasons (C). Crop use efficiency, as defined as grape yield per mm of irrigation applied by vine, for 2019 (D), 2020 (E), and 2021 (F). Heatwave events (3 or more days predicted to have a maximum temperature above the 38 degrees Celsius threshold) are highlighted in gray. Error bars are SEM (n=27). P < 0.05; significance was evaluated by one-way analysis of variance and Tukey HSD was used to study post hoc differences among treatments, letters denote significant differences.

**Figure 5.**
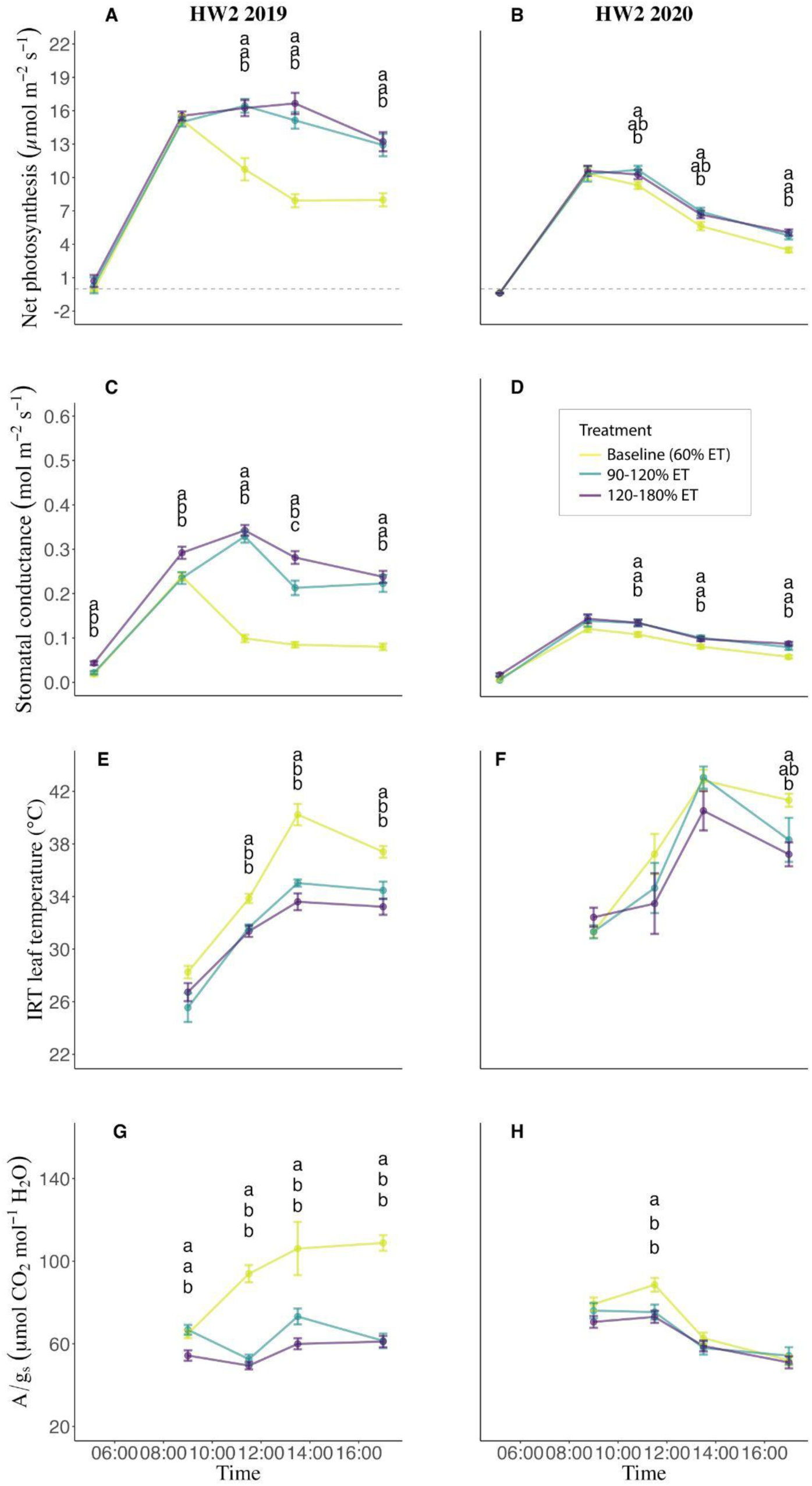
Diurnal measurements of transpiration, E (top) instantaneous WUE, A/E (middle), and intrinsic WUE, A/g_s_ (bottom) carried out during HW2_2019_ (A, C, E) and HW2_2020_ (B, D, F) on August 15^th^, 2019 and July 12^th^, 2020, respectively. Error bars are SEM (n=9 for predawn measurements and for the rest of diurnal measurements n=27). P < 0.05; significance was evaluated by one-way analysis of variance and Tukey HSD was used to study post hoc differences among treatments, letters denote significant differences.

### Yield components

Cluster weight, total yield per vine, and grape berry weight were components of yield recorded at harvest across all treatments. The baseline treatment had significantly lower yield per vine than the supplemental irrigation treatments across all years. In 2019, this was in accordance with a significant decrease in berry and cluster weight. In 2020 and 2021, only cluster weight was lower, albeit non-significant in 2021. There was no significant difference in berry weight in 2020 or 2021. Across all years, the moderate irrigation treatment (90-120% ET) produced the highest vine yield, and importantly there was a decrease in mean yield with the highest irrigation treatment across all years.

### Berry composition

Total soluble solids (or °Brix), pH, and titratable acidity (TA), all measures used to determine harvest timing in grapes, were all significantly impacted by HW irrigation regime in two of the three years of the experiment (Table 1). In 2019, no significant differences were seen in TSS due to variable harvest times, however, higher values persisted through the week before harvest in the baseline relative to the supplemental irrigation treatments (Table S18). At harvest, TSS in 2020 was significantly higher in the baseline, while in 2021, there were no significant differences between treatments (Table 1). In 2019, the 120% ET treatment had grapes with significantly lower pH and higher TA than the baseline and 180% ET treatments, but these differences were not present at harvest. In contrast, the 2020 baseline treatment yielded fruit with high pH and lower TA relative to both supplemental irrigation treatments. In 2021, there were no significant treatment differences in pH, but the TA was significantly lower in the baseline treatment at harvest (Table 1).

**Table 1.** Yield components and primary chemistry of the 2019, 2020 and 2021 growing seasons taken at harvest of vines exposed to differential irrigation prior to and during HWs.

| 2019 |  |  |  |  |
| --- | --- | --- | --- | --- |
|  | Baseline (60%ET) | 2x Baseline ET | 3x Baseline ET | P-value |
| Brix | 25.62 ± 0.1<sup>a</sup> | 25.79 ± 0.17<sup>a</sup> | 25.51 ± 0.14<sup>a</sup> | ns |
| pH | 3.67 ± 0.02<sup>a</sup> | 3.5 ± 0.03<sup>b</sup> | 3.66 ± 0.01<sup>a</sup> | *** |
| Titrateable acidity (g/L) | 3.91 ± 0.09<sup>b</sup> | 4.95 ± 0.16<sup>a</sup> | 4.27 ± 0.06<sup>b</sup> | *** |
| Yield (kg/plant) | 6.25 ± 0.23<sup>b</sup> | 8.86 ± 0.63<sup>a</sup> | 7.7 ± 0.58<sup>ab</sup> | * |
| Weight (gr/cluster) | 67.27 ± 2.87<sup>b</sup> | 83.52 ± 4.63<sup>a</sup> | 78.59 ± 3.78<sup>ab</sup> | * |
| Berry weight (gr) | 0.89±0.01<sup>b</sup> | 1.05±0.02<sup>a</sup> | 1.03±0.01<sup>a</sup> | *** |
| 2020 |  |  |  |  |
|  | Baseline (60%ET) | 2x Baseline ET | 3x Baseline ET | P-value |
| Brix | 26.88 ± 0.16<sup>a</sup> | 25.89 ± 0.23<sup>b</sup> | 26 ± 0.22<sup>b</sup> | ** |
| pH | 3.66 ± 0.01<sup>a</sup> | 3.54 ± 0.02<sup>b</sup> | 3.54 ± 0.02<sup>b</sup> | *** |
| Titrateable acidity (g/L) | 4.02 ± 0.08<sup>b</sup> | 4.54 ± 0.13<sup>a</sup> | 4.86 ± 0.12<sup>a</sup> | *** |
| Yield (kg/plant) | 7.28 ± 0.17<sup>b</sup> | 8.32 ± 0.2<sup>a</sup> | 8.24 ± 0.2<sup>a</sup> | *** |
| Weight (gr/cluster) | 75.72 ± 1.81<sup>b</sup> | 82.72 ± 1.96<sup>a</sup> | 80.99 ± 1.63<sup>ab</sup> | * |
| Berry weight (gr) | 0.95 ± 0.01<sup>a</sup> | 0.98 ± 0.01<sup>a</sup> | 0.97 ± 0.01<sup>a</sup> | ns |
| 2021 |  |  |  |  |
|  | Baseline (60%ET) | 1.5x Baseline ET | 2x Baseline ET | P-value |
| Brix | 24.58 ± 0.19<sup>a</sup> | 24.53 ± 0.18<sup>a</sup> | 24.66 ± 0.25<sup>a</sup> | ns |
| pH | 3.75 ± 0.02<sup>a</sup> | 3.69 ± 0.02<sup>a</sup> | 3.72 ± 0.02<sup>a</sup> | ns |
| Titrateable acidity (g/L) | 3.2 ± 0.05<sup>b</sup> | 3.42 ± 0.05<sup>a</sup> | 3.42 ± 0.04<sup>a</sup> | ** |
| Yield (kg/plant) | 7.4 ± 0.27<sup>b</sup> | 8.89 ± 0.29<sup>a</sup> | 8.33 ± 0.27<sup>ab</sup> | *** |
| Weight (gr/cluster) | 80.94 ± 3.56<sup>a</sup> | 85.78 ± 3.02<sup>a</sup> | 76.41 ± 2.36<sup>a</sup> | ns |
| Berry weight (gr) | 1.06 ± 0.01<sup>a</sup> | 1.09 ± 0.01<sup>a</sup> | 1.04 ± 0.02<sup>a</sup> | ns |

\*, \*\*, and \*\*\* for P values <0.05, <0.01 and <0.001 of one-way ANOVA.
Different letters denote significant differences among treatments for each variable studied after performing a Tukey HSD test (p<0.05).

## Discussion

As heat extremes and water availability become increasingly problematic for cropping systems, it will be critical to understand how plant available water and the resulting irrigation decisions during heat wave events impact plant physiological performance and productivity. Higher temperatures, periods of drought, and associated rises in vapor pressure deficit, which result in reductions in stomatal conductance, leaf cooling ability, and decreases in carbon uptake, will dramatically impact plant yields critical to agricultural productivity (Grossiord *et al*., 2020; Koehler *et al*., 2023; Novick *et al*., 2024).

Strategic irrigation timing is critical for perennial crops during heat events, as water application is the primary strategy employed to mitigate heat stress responses in irrigated agriculture systems (Table S1, (Parker *et al*., 2020). Deficit irrigation has emerged as a promising approach to both save water and induce beneficial plant physiological regulations, including optimized stomatal opening and balanced reproductive and vegetative growth (Chaves *et al*., 2010; Keller *et al*., 2016). Irrigation management aims to increase water use efficiency (WUE), which can be improved from the stomatal to the regional scale through strategic water application (Davies *et al*., 2002). However, excessive irrigation can promote excessive vegetative growth with negative impacts on berry quality, including reduced color and sugar content, while also increasing disease incidence through larger canopy leaf area (Bravdo *et al*., 1984; Dokoozlian & Kliewer, 1995; Costa *et al*., 2007; Chaves *et al*., 2010). The treatments used in our current study represent the realistic range used in commercial grape growing that could result in detrimental stress or overwatering in response to HWs. The results of our study, where we applied variable amounts of irrigation during extreme heat events over three growing seasons, show that applying supplemental irrigation before and during heatwaves is critical to maintaining physiological function, production and crop water use efficiency of the vines, but that excess water (above 2X a baseline ET of 60%) did not provide any additional benefit and resulted in a reduction in crop WUE, plant productivity, and fruit quality. These findings align with previous work demonstrating that deficit irrigation can improve WUE while controlling vigor and production quality when appropriately managed (Costa *et al*., 2007), but that excessive water can undermine these benefits.

### Water Status Responses to Differential Irrigation

The baseline treatment showed variability in water deficit intensity and HW responses across years, ranging from mild water deficit in 2019 to greater deficit post-veraison in 2020, and minimal differences in 2021 (Figs. 2 & S2). These patterns differ from previous studies in Shiraz grapevines with continuous temperature and irrigation treatments (Sadras *et al*., 2012; Bonada *et al*., 2013, 2015), likely because our experiment involved natural heatwaves with irrigation applied only one to two days before and during the heat events. Despite large differences in water application between the supplemental irrigation treatments (90-180% ET), no significant differences were found between these treatments for midday Ψ_stem_ measurements across three years (Fig. S2), suggesting that irrigation in the 90-120% ET treatment was sufficient for complete recovery of water loss through evapotranspiration. Further, under high VPD conditions, midday water potential measurements most typically used in management of perennial agriculture(Choné, 2001; Shackel, 2007; Acevedo-Opazo *et al*., 2010) may not suffice as reliable or telling indicators of stress or the need to for mitigation measures, as pre-dawns and measures of conductance can be more informative in assessing potential impacts of heat and drought stress on yield and quality.

### Gas exchange responses before and after heatwave events

Gas exchange responses followed consistent patterns across all three years, with significant reductions in net photosynthesis and stomatal conductance during HWs that persisted 4-5 days post-HW (Fig. 3A-F). However, the baseline treatment showed evident physiological recovery during subsequent sampling dates. The reduction in carbon assimilation in the baseline treatment likely reflects stomatal limitation and gas exchange regulation driven by thermal and water stress. Previous studies report similar photosynthetic rate decreases during transient heat events (Gouot *et al*., 2019b; Greer & Weston, 2010; Greer & Weedon, 2013), though responses vary with environmental conditions and genotype. Some studies attributed reductions to stomatal closure (Greer & Weston, 2010), while others found biochemical limitations in the electron transport chain and Rubisco activity during longer heat waves (Greer & Weedon, 2013). Conversely, well-watered Shiraz has shown upregulated gas exchange during short heat waves (Soar *et al*., 2009), demonstrating genotypic plasticity for increasing transpiration and achieving canopy cooling. The baseline treatment consistently exhibited lower transpiration rates during and post-HWs across three years, generally aligning with higher leaf temperatures, though differences were not always significant. Higher leaf temperatures in the baseline indicate decreased transpiration due to stomatal limitation from higher atmospheric demand and lower water availability. Importantly, the similar physiological performance of the 90-120% ET and 120-180% ET treatments (Fig. 3) indicates no additional cooling benefit beyond 100% ET recovery.

While stomatal closure is typically coupled with lower water potential under drought (Comstock, 2002), the significant reduction in stomatal conductance in the baseline treatment during 2019 heatwaves did not align with midday Ψ_stem_ and Ψ_leaf_ responses (Fig. 2A, 3D, Fig. S2). However, the baseline showed significantly lower post-HW water potentials compared to other treatments, confirming that differential irrigation affected plant water status as expected. Similar decoupling occurred in 2020 and 2021, with stomatal responses showing variable relationships to water potential across heat events (Fig. 2B,C; Fig. 3E,F; Fig. S2). These results highlight a lack of clear correlation between plant water status and stomatal responses under extreme heat, which is particularly important for woody perennial crops where Ψ_stem_ typically guides irrigation management (Choné, 2001; Deloire & Heyns, 2011). The variable physiological responses observed across the three years and the consistent post-HW recovery in the baseline treatment suggest that acclimation may play an important role in grapevine heat stress responses. Previously heat-stressed vines can show faster recovery than non-acclimated vines (Carvalho *et al*., 2015), and plants possess stress memory mechanisms that allow them to respond more effectively to recurring stresses (Bruce *et al*., 2007; Crisp *et al*., 2016; Staacke *et al*., 2025). Within-season acclimation occurs through rapid adjustments in gene expression, metabolite accumulation, and physiological processes, while across-season acclimation may involve epigenetic modifications and structural changes that persist between growing seasons (Lämke & Bäurle, 2017). The baseline vines in our study experienced repeated heat stress across three consecutive seasons, potentially developing enhanced stress tolerance through priming effects. This could explain the evident physiological recovery observed 4-5 days post-HW despite lower water availability and greater stress during events. However, the yield reductions observed across all three years suggest that while short-term physiological recovery occurred, the cumulative effects of repeated stress still impacted reproductive development and productivity. The threshold between beneficial acclimation and detrimental cumulative stress likely depends on stress intensity, duration, frequency, and the recovery period between events (Mittler, 2006; Suzuki *et al*., 2014).

### Leaf-level and crop water use efficiency

An important distinction emerged between leaf-level and crop-level water use efficiency in our study. The baseline treatment, experiencing greater drought and heat stress, consistently showed higher instantaneous leaf-level WUE across all three years, reflecting the expected physiological response, where stomatal closure reduces transpiration more proportionally than photosynthesis, thereby improving the ratio of carbon gain to water loss (Chaves *et al*., 2010; Medrano *et al*., 2015). Despite this improvement in instantaneous efficiency, the 90-120% ET treatment achieved higher crop-level WUE all years (Figures 4D-I, S1-S3). This apparent paradox highlights the trade-off inherent in deficit irrigation strategies: while stressed vines conserve water through stomatal regulation and achieve higher momentary efficiency, their reduced total carbon assimilation and lower yield result in poorer overall water-to-productivity ratios at the whole-plant and crop level (Flexas *et al*., 2010). The 90-120% ET treatment’s superior crop-level WUE, coupled with highest yields and maintained fruit quality (Campbell et al. 2025, in revision), demonstrates that moderate supplemental irrigation during heat waves optimizes the balance between water conservation and agricultural productivity, a critical consideration for sustainable viticulture and woody perennial crop production in water-scarce regions facing increasing heat stress.

### Yield and Berry Compositional Differences at Harvest

Across three consecutive years (2019-2021), the baseline irrigation treatment consistently showed significantly lower yields and berry weights compared to higher irrigation treatments. Multiple factors contributed to these yield reductions. Water availability and temperature strongly impact both current and subsequent season inflorescence formation and yield (Carmo Vasconcelos *et al*., 2009; Guilpart *et al*., 2014; Pagay & Collins, 2017). The 2020 baseline yield decrease likely reflected reduced fruit set and ovule fertility from heat stress during flowering, not solely berry weight reduction—particularly since baseline cluster weight was lowest, though only 120% ET was significantly higher. The 2021 yield decrease may represent a carryover effect from the 2020 flowering-period heatwave (Pagay & Collins, 2017).

Post-fruit set berry weight decreases from transient heat exposure and water deficit are well-documented (Roby & Matthews, 2004; Roby *et al*., 2004; Soar *et al*., 2009; Casassa *et al*., 2015). Phase I grape development involves intensive cell division (Coombe and Iland 2004) that is sensitive to both heat (Kliewer, 1977) and water stress, with (Bonada *et al*., 2015) demonstrating additive effects leading to decreased berry size. Following post-veraison heatwaves (HW2_2019_ and HW3_2020_), the baseline experienced further weight decreases (Fig. S5), consistent with research showing that high temperatures cause cell expansion cessation during veraison and mid-ripening (Greer & Weedon, 2013). Increased berry dehydration and shriveling after post-veraison heatwaves may have also contributed to lower baseline berry weight (Caravia *et al*., 2016; Martínez-Lüscher *et al*., 2020), though shriveling percentage was not assessed. Despite three years of persistent yield differences, no treatment differences in carbohydrate reserves across different organs were found at study end (Furze et al. 2025, in prep), suggesting carbohydrate depletion did not occur.

Significant treatment effects and heatwave-driven TSS changes occurred across all three years, both pre- and post-veraison. Paradoxically, the baseline treatment reached significantly higher Brix during or after heat events (Tables S18-S20), despite showing declined stomatal conductance and decreased photosynthesis during heatwaves (Fig. 3A-F), which would suggest lower carbon availability for hexose accumulation and consequently delayed ripening. Several factors explain this apparent contradiction. The consistent baseline berry weight and yield declines would concentrate hexoses, increasing TSS. Post-heatwave gas exchange measurements showed physiological stress recovery in the baseline, indicating only short, transient carbon assimilation limitation. Additionally, lower water availability during heatwaves may have stimulated ripening through abscisic acid stress signaling (Castellarin *et al*., 2007a)). This contrasts with findings by (Soar *et al*., 2009), who observed maintained gas exchange without changes in berry size or sugar accumulation in well-irrigated Shiraz vines exposed to three consecutive heat days, demonstrating genotypic plasticity to adapt under challenging conditions (Coupel-Ledru *et al*., 2024).

The baseline treatment consistently showed lower TA coupled with higher pH across all years (Table 1), consistent with previous findings that elevated post-veraison temperatures and deficit irrigation promote higher respiration and malic acid consumption (Coombe, 1987; Keller *et al*., 2008; Shellie, 2011; Sweetman *et al*., 2014; Casassa *et al*., 2015; Bonada *et al*., 2015; Pastore *et al*., 2017). These results are relevant to wine making, as lowering acidity to achieve desired quality can result in costly modification in the winery. Interestingly, the 180% ET treatment showed significantly higher pH compared to 120% ET in 2019 and significantly lower TA (Table 1), potentially due to a dilution effect from increased berry size decreasing organic acid concentration. However, 120% ET had the highest berry weight despite no significant differences between 120% ET and 180% ET at harvest. Overall, the 90-120% ET treatment showed optimal results in physiological responses and grape quality traits for both primary and secondary metabolites (Campbell et al. 2025, in revision), suggesting moderate supplemental irrigation optimizes the balance between sustainable water use and crop performance.

While most previous work has focused on altering irrigation and heat for longer periods of time or under artificial conditions (e.g., chamber or pot experiments), our study was more realistic in that natural heat events necessitate altering irrigation or increasing water availability episodically, or during the timing of the heat event. Climatic chamber experiments with grapevine are relatively complicated and costly due to the perennial nature and annual reproductive cycle of the vine, resulting in most experiments being conducted in the field where fine temperature control is impossible but water availability can be managed. Field conditions introduce biases from environmental fluctuations that are difficult to circumvent but reflect the reality of transient variations in light, air movement, and moisture that affect stomatal conductance, plant surface temperature, and physiological processes in commercial settings. They also allow for more realistic extension to other species and crops. Moreover, controlled environment studies typically apply continuous heat treatments (e.g., (Carvalho *et al*., 2015)), while field-grown vines experience naturally variable temperature regimes where heat events may be intermittent and of varying duration (Greer & Weedon, 2012, 2013). Previous studies have also recognized that the morpho-anatomical and physiological traits involved in heat stress tolerance vary with genotype-environment interactions, seasonal changes, stress intensity, and duration (Bruce *et al*., 2007; Costa *et al*., 2012; Ahrens *et al*., 2021; Grossman, 2023; Zhou *et al*., 2023), emphasizing the importance of studying responses under realistic field conditions.

Our results show that altering irrigation only during heat events impacts plant physiological responses, and downstream crop yield and quality. This approach contrasts with studies applying continuous deficit irrigation strategies throughout the season (Chaves *et al*., 2010; Costa *et al*., 2012), but better reflects the practical management decisions growers must make when heat waves occur (Romero *et al*., 2022). Event-based irrigation management is particularly relevant given that severe deficit irrigation should be avoided before heat waves to prevent vines from being excessively stressed during the event, while supplemental irrigation applied during or immediately after heat stress can help restore vine water status and maintain gas exchange. Avoiding severe water deficits before heat events and applying targeted supplemental irrigation during or immediately after stress can optimize water status and sustain photosynthesis, highlighting the value of physiology-informed irrigation management under increasingly frequent and intense heat events.

## Supporting information

Survey Results Summary

## Acknowledgements

Support from E. & J. Gallo and their staff was critical to the success of the field trial, seed funding from USDA Climate Hub, CDFA Specialty Crop Block Grant Award No. 19-0001-013-SF, Department of Viticulture & Enology and College of Environmental and Agricultural Science at UC Davis.

## Author Contributions

EJF, MG and AJM planned and designed the research, wrote manuscript. LS and ND supported and helped plan research. MG, SB, KE, and EJF conducted field and lab work. NB helped design research, provided feedback and edits. LP and EJF planned and conducted grower survey. EJF and MG performed data analysis and interpretation. All authors edited and approved manuscript.

## References

Abatzoglou JT, Brown TJ. 2012. A comparison of statistical downscaling methods suited for wildfire applications. International Journal of Climatology 32: 772–780.

Acevedo-Opazo C, Ortega-Farias S, Fuentes S. 2010. Effects of grapevine (Vitis vinifera L.) water status on water consumption, vegetative growth and grape quality: An irrigation scheduling application to achieve regulated deficit irrigation. Agricultural Water Management 97: 956–964.

Ahrens CW, Challis A, Byrne M, Leigh A, Nicotra AB, Tissue D, Rymer P. 2021. Repeated extreme heatwaves result in higher leaf thermal tolerances and greater safety margins. The New Phytologist 232: 1212–1225.

Albano CM, Abatzoglou JT, McEvoy DJ, Huntington JL, Morton CG, Dettinger MD, Ott TJ. 2022. A Multidataset Assessment of Climatic Drivers and Uncertainties of Recent Trends in Evaporative Demand across the Continental United States. Journal of Hydrometeorology 23: 505–519.

Allen RG, Pereira LS, Raes D, Smith M. 1998. Crop evapotranspiration: Guidelines for computing crop water requirements. Rome, Italy: Food and Agriculture Organization of the United Nations.

Bedsworth L, Cayan D, Franco G, Fisher L, Ziaja S. 2018. California’s fourth climate change assessment: Statewide summary report. Publication number: SUM-CCCA4-2018-013.

Bonada M, Jeffery DW, Petrie PR, Moran MA, Sadras VO. 2015. Impact of elevated temperature and water deficit on the chemical and sensory profiles of Barossa Shiraz grapes and wines: Temperature and water effects on grapes and wines. Australian journal of grape and wine research 21: 240–253.

Bonada M, Sadras V, Moran M, Fuentes S. 2013. Elevated temperature and water stress accelerate mesocarp cell death and shrivelling, and decouple sensory traits in Shiraz berries. Irrigation Science 31: 1317–1331.

Bravdo B, Hepner Y, Loinger C, Cohen S, Tabacman H. 1984. Effect of crop level on growth, yield and wine quality of a high yielding carignane vineyard. American Journal of Enology and Viticulture 35: 247–252.

Breshears DD, Fontaine JB, Ruthrof KX, Field JP, Feng X, Burger JR, Law DJ, Kala J, Hardy GESJ. 2021. Underappreciated plant vulnerabilities to heat waves. The New phytologist 231: 32–39.

Bruce TJA, Matthes MC, Napier JA, Pickett JA. 2007. Stressful ‘memories’ of plants: Evidence and possible mechanisms. Plant Science: An International Journal of Experimental Plant Biology 173: 603–608.

Campbell JR, Galeano M, McElrone AJ, Sanchez L, Dookozlian N, Bagshaw S, Waterhouse, AL, Forrestel EJ. 2026. Enhanced Irrigation during Extreme Heat Events Preserves Anthocyanins in Cabernet Sauvignon. Journal of Agriculture & Food Chemistry. In revision.

Caravia L, Collins C, Petrie PR, Tyerman SD. 2016. Application of shade treatments during Shiraz berry ripening to reduce the impact of high temperature: Shade reduces impact of high temperature on Shiraz. Australian journal of grape and wine research 22: 422–437.

Carmo Vasconcelos M, Greven M, Winefield CS, Trought MCT, Raw V. 2009. The Flowering Process of Vitis vinifera: A Review. American journal of enology and viticulture 60: 411–434.

Carvalho LC, Coito JL, Colaço S, Sangiogo M, Amâncio S. 2015. Heat stress in grapevine: the pros and cons of acclimation. Plant, cell & environment 38: 777–789.

Casassa LF, Keller M, Harbertson JF. 2015. Regulated deficit irrigation alters anthocyanins, tannins and sensory properties of Cabernet Sauvignon grapes and wines. Molecules 20: 7820–7844.

Castellarin SD, Matthews MA, Di Gaspero G, Gambetta GA. 2007a. Water deficits accelerate ripening and induce changes in gene expression regulating flavonoid biosynthesis in grape berries. Planta 227: 101–112.

Chaves MM, Zarrouk O, Francisco R, Costa JM, Santos T, Regalado AP, Rodrigues ML, Lopes CM. 2010. Grapevine under deficit irrigation: hints from physiological and molecular data. Annals of botany 105: 661–676.

Choné X. 2001. Stem water potential is a sensitive indicator of grapevine water status. Annals of Botany 87: 477–483.

Comstock JP. 2002. Hydraulic and chemical signalling in the control of stomatal conductance and transpiration. Journal of experimental botany 53: 195–200.

Coombe BG. 1987. Influence of Temperature on Composition and Quality of Grapes. Acta horticulturae 206: 23–36.

Coombe B, Iland P. 2004. Grape Berry Development and Winegrape Quality. *In ‘*Viticulture. Vol. 1. Resources*’. (Eds* PR Dry, BG Coombe): 210–248.

Costa JM, Ortuño MF, Chaves MM. 2007. Deficit irrigation as a strategy to save water: Physiology and potential application to horticulture. Journal of integrative plant biology 49: 1421–1434.

Costa JM, Ortu O MF, Lopes CM, Chaves MM. 2012. Grapevine varieties exhibiting differences in stomatal response to water deficit. Functional plant biology: FPB 39: 179–189.

Coupel-Ledru A, Westgeest AJ, Albasha R, Millan M, Pallas B, Doligez A, Flutre T, Segura V, This P, Torregrosa L, et al. 2024. Clusters of grapevine genes for a burning world. The New phytologist 242: 10–18.

Crisp PA, Ganguly D, Eichten SR, Borevitz JO, Pogson BJ. 2016. Reconsidering plant memory: Intersections between stress recovery, RNA turnover, and epigenetics. Science Advances 2: e1501340.

Davies WJ, Wilkinson S, Loveys B. 2002. Stomatal control by chemical signalling and the exploitation of this mechanism to increase water use efficiency in agriculture. The New Phytologist 153: 449–460.

Deloire A, Heyns D. 2011. The leaf water potentials: Principles, method and thresholds. Wynboer 265: 119–121.

Diffenbaugh NS, Davenport FV, Burke M. 2021. Historical warming has increased U.S. crop insurance losses. Environmental research letters: ERL [Web site] 16: 084025.

Dokoozlian NK, Kliewer WM. 1995. The light environment within grapevine canopies. II. Influence of leaf area density on fruit zone light environment and some canopy assessment parameters. American Journal of Enology and Viticulture 46: 219–226.

Fang Z, Zhang W, Brandt M, Abdi AM, Fensholt R. 2022. Globally increasing atmospheric aridity over the 21st century. Earth’s future 10.

Flexas J, Galmãs J, Gallã A, Gulãas J, Pou A, Ribas-Carbo M, Tomãs M, Medrano H. 2010. Improving water use efficiency in grapevines: potential physiological targets for biotechnological improvement. Australian journal of grape and wine research 16: 106–121.

Furze FE, Rodriguez-Urquidi A, Galeano M, Lozano J, Sanchez L, Dookozlian N, McElrone AJ, Forrestel EJ. Carbohydrate reserves recover in wine grapes regardless of irrigation manipulation during heat waves. In review.

Gershunov A, Cayan DR, Retornaz B. 2010. California Heat Waves with Impacts on Wine Grapes. The Ocean, the Wine, and the Valley: The Lives of Antoine Badan, EG Pavia, J. Sheinbaum and J. Candela (Eds.): 205–224.

Gershunov A, Guirguis K. 2012. California heat waves in the present and future. Geophysical research letters 39: L18710.

Gouot J, Smith J, Holzapfel B, Barril C. 2019b. Single and cumulative effects of whole-vine heat events on Shiraz berry composition. OENO One 53: 171–187.

Greer DH, Weedon MM. 2012. Modelling photosynthetic responses to temperature of grapevine (Vitis vinifera cv. Semillon) leaves on vines grown in a hot climate. Plant, cell & environment 35: 1050–1064.

Greer DH, Weedon MM. 2013. The impact of high temperatures on Vitis vinifera cv. Semillon grapevine performance and berry ripening. Frontiers in plant science 4: 491.

Greer D, Weston C. 2010. Heat stress affects flowering, berry growth, sugar accumulation and photosynthesis of Vitis vinifera cv. Semillon grapevines grown in a controlled environment. Functional plant biology: FPB 37: 206–214.

Grossiord C, Buckley TN, Cernusak LA, Novick KA, Poulter B, Siegwolf RTW, Sperry JS, McDowell NG. 2020. Plant responses to rising vapor pressure deficit. The New phytologist 226: 1550–1566.

Grossman JJ. 2023. Phenological physiology: seasonal patterns of plant stress tolerance in a changing climate. The New Phytologist 237: 1508–1524.

Guilpart N, Metay A, Gary C. 2014. Grapevine bud fertility and number of berries per bunch are determined by water and nitrogen stress around flowering in the previous year. European journal of agronomy: the journal of the European Society for Agronomy 54: 9–20.

Iland P, Bruer N, Edwards G, Weeks S, Wilkes E. 2004. Chemical analysis of grapes and wine: techniques and concepts.,(Patrick Iland Wine Promotions PTY Ltd: Campbelltown, SA). New South Wales, Australia.

Jones GV, Davis RE. 2000. Climate influences on grapevine phenology, grape composition, and wine production and quality for Bordeaux, France. American Journal of Enology and Viticulture 51: 249–261.

Jones GV, White MA, Cooper OR, Storchmann K. 2005. Climate change and global wine quality. Climatic change 73: 319–343.

Keller M. 2010. Managing grapevines to optimise fruit development in a challenging environment: a climate change primer for viticulturists. Australian journal of grape and wine research 16: 56–69.

Keller M, Romero P, Gohil H, Smithyman RP, Riley WR, Federico Casassa L, Harbertson JF. 2016. Deficit Irrigation Alters Grapevine Growth, Physiology, and Fruit Microclimate. American journal of enology and viticulture 67: 426–435.

Keller M, Smithyman RP, Mills LJ. 2008. Interactive Effects of Deficit Irrigation and Crop Load on Cabernet Sauvignon in an Arid Climate. American journal of enology and viticulture 59: 221–234.

Kliewer WM. 1977. Effect of High Temperatures during the Bloom-Set Period on Fruit-Set, Ovule Fertility, and Berry Growth of Several Grape Cultivars. American journal of enology and viticulture 28: 215–222.

Koehler T, Wankmüller FJP, Sadok W, Carminati A. 2023. Transpiration response to soil drying versus increasing vapor pressure deficit in crops: physical and physiological mechanisms and key plant traits. Journal of Experimental Botany 74: 4789–4807.

Kool D, Agam N, Lazarovitch N, Heitman JL, Sauer TJ, Ben-Gal A. 2014. A review of approaches for evapotranspiration partitioning. Agricultural and forest meteorology 184: 56–70.

Kriedemann PE. 1968. Photosynthesis in vine leaves as a function of light intensity, temperature, and leaf age. Vitis 7: 213–220.

Kustas WP, McElrone AJ, Agam N, Knipper K. 2022. From vine to vineyard: the GRAPEX multi-scale remote sensing experiment for improving vineyard irrigation management. Irrigation Science 40: 435–444.

Lämke J, Bäurle I. 2017. Epigenetic and chromatin-based mechanisms in environmental stress adaptation and stress memory in plants. Genome Biology 18: 124.

Lee H, Calvin K, Dasgupta D, Krinner G, Mukherji A, Thorne P, Trisos C, Romero J, Aldunce P, Barrett K, et al. 2023. Climate change 2023: synthesis report. Contribution of working groups I, II and III to the sixth assessment report of the intergovernmental panel on climate change. The Australian National University.

van Leeuwen C, Sgubin G, Bois B, Ollat N, Swingedouw D, Zito S, Gambetta GA. 2024. Climate change impacts and adaptations of wine production. Nature Reviews Earth & Environment 5: 258–275.

Manabe S, Wetherald RT, Milly PCD, Delworth TL, Stouffer RJ. 2004. Century-scale change in water availability: CO 2-quadrupling experiment. Climatic change 64: 59–76.

Martínez-Lüscher J, Kurtural S, Chen C, Brillante L. 2020. Fruit Zone Overexposure and Heat Events Lead to Severe Losses of Anthocyanins and Marketable Crop in ‘Cabernet Sauvignon’ Wine Grape under Two Irrigation Amounts. The Annals of applied biology.

Medrano H, Tomás M, Martorell S, Escalona J-M, Pou A, Fuentes S, Flexas J, Bota J. 2015. Improving water use efficiency of vineyards in semi-arid regions. A review. Agronomy for sustainable development 35: 499–517.

Mittler R. 2006. Abiotic stress, the field environment and stress combination. Trends in Plant Science 11: 15–19.

Morales-Castilla I, García de Cortázar-Atauri I, Cook BI, Lacombe T, Parker A, van Leeuwen C, Nicholas KA, Wolkovich EM. 2020. Diversity buffers winegrowing regions from climate change losses. Proceedings of the National Academy of Sciences of the United States of America 117: 2864–2869.

Novick KA, Ficklin DL, Grossiord C, Konings AG, Martínez-Vilalta J, Sadok W, Trugman AT, Williams AP, Wright AJ, Abatzoglou JT, et al. 2024. The impacts of rising vapour pressure deficit in natural and managed ecosystems. Plant, Cell & Environment 47: 3561–3589.

Pagay V, Collins C. 2017. Effects of timing and intensity of elevated temperatures on reproductive development of field-grown Shiraz grapevines. OENO One 51.

Parker LE, McElrone AJ, Ostoja SM, Forrestel EJ. 2020. Extreme heat effects on perennial crops and strategies for sustaining future production. Plant science: an international journal of experimental plant biology: 110397.

Parker LE, Zhang N, Abatzoglou JT, Kisekka I, McElrone AJ, Ostoja SM. 2024. A variety-specific analysis of climate change effects on California winegrapes. International journal of biometeorology.

Pastore C, Dal Santo S, Zenoni S, Movahed N, Allegro G, Valentini G, Filippetti I, Tornielli GB. 2017. Whole Plant Temperature Manipulation Affects Flavonoid Metabolism and the Transcriptome of Grapevine Berries. Frontiers in plant science 8: 929.

Qin Y, Abatzoglou JT, Siebert S, Huning LS, AghaKouchak A, Mankin JS, Hong C, Tong D, Davis SJ, Mueller ND. 2020. Agricultural risks from changing snowmelt. Nature climate change 10: 459–465.

Robinson PJ. 2001. On the definition of a heat wave. Journal of Applied Meteorology 40: 762–775.

Roby G, Harbertson JF, Adams DA, Matthews MA. 2004. Berry size and vine water deficits as factors in winegrape composition: Anthocyanins and tannins. Australian journal of grape and wine research 10: 100–107.

Roby G, Matthews MA. 2004. Relative proportions of seed, skin and flesh, in ripe berries from Cabernet Sauvignon grapevines grown in a vineyard either well irrigated or under water deficit. Australian journal of grape and wine research 10: 74–82.

Romero P, Navarro JM, Ordaz PB. 2022. Towards a sustainable viticulture: The combination of deficit irrigation strategies and agroecological practices in Mediterranean vineyards. A review and update. Agricultural water management 259: 107216.

Sadras VO, Montoro A, Moran MA, Aphalo PJ. 2012. Elevated temperature altered the reaction norms of stomatal conductance in field-grown grapevine. Agricultural and Forest Meteorology 165: 35–42.

Sadras VO, Moran MA. 2013. Asymmetric warming effect on the yield and source:sink ratio of field-grown grapevine. Agricultural and Forest Meteorology 173: 116–126.

Sanchez LA, Sams B, Alsina MM, Hinds N, Klein LJ, Dokoozlian N. 2017. Improving vineyard water use efficiency and yield with variable rate irrigation in California. Advances in Animal Biosciences 8: 574–577.

Sgubin G, Swingedouw D, Mignot J, Gambetta GA, Bois B, Loukos H, Noël T, Pieri P, García de Cortázar-Atauri I, Ollat N, et al. 2023. Non-linear loss of suitable wine regions over Europe in response to increasing global warming. Global change biology 29: 808–826.

Shackel KA. 2007. Water relations of woody perennial plant species. OENO One 41: 121.

Shellie KC. 2011. Interactive Effects of Deficit Irrigation and Berry Exposure Aspect on Merlot and Cabernet Sauvignon in an Arid Climate. American journal of enology and viticulture 62: 462–470.

Soar CJ, Collins MJ, Sadras VO. 2009. Irrigated Shiraz vines (Vitis vinifera) upregulate gas exchange and maintain berry growth in response to short spells of high maximum temperature in the field. Functional Plant Biology 36: 801–814.

Staacke T, Mueller-Roeber B, Balazadeh S. 2025. Stress resilience in plants: the complex interplay between heat stress memory and resetting. The New Phytologist 245: 2402–2421.

Suzuki N, Rivero RM, Shulaev V, Blumwald E, Mittler R. 2014. Abiotic and biotic stress combinations. The New Phytologist 203: 32–43.

Sweetman C, Sadras VO, Hancock RD, Soole KL, Ford CM. 2014. Metabolic effects of elevated temperature on organic acid degradation in ripening Vitis vinifera fruit. Journal of experimental botany 65: 5975–5988.

White MA, Diffenbaugh NS, Jones GV, Pal JS, Giorgi F. 2006. Extreme heat reduces and shifts United States premium wine production in the 21st century. Proceedings of the National Academy of Sciences of the United States of America 103: 11217–11222.

Yin C, Yang Y, Chen X, Yue X, Liu Y, Xin Y. 2022. Changes in global heat waves and its socioeconomic exposure in a warmer future. Climate Risk Management 38: 100459.

Zhou C, Wu S, Li C, Quan W, Wang A. 2023. Response mechanisms of woody plants to high-temperature stress. Plants 12: 3643.

