## Supplementary material for "Optimizing irrigation during heat events sustains grapevine physiology and fruit production": 1_SupplementalLegends

**Supplemental Figures & Tables**

**Figure S1.** Boxplot (i.e., minimum, 25th percentile, median, 75th percentile, and maximum) for WUE at the crop level of treatments exposed to differential irrigation (Baseline (60% ET), 90% ET, and 120% ET) at harvest in 2019 (n=3).

**Figure S2.** Predawn (A-C) and midday (D-F) leaf water potential measurements and significance based for 2019 through 2021. Values are means ± SE (n=9). P < 0.05; significance was evaluated by one-way analysis of variance and Tukey HSD was used to study post hoc differences among treatments, letters denote significant differences.

**Table S1.** Survey results.

**Table S2.** Midday stem water potentials (Ψ _stem_) from vines with three differential irrigation treatments during HWs, Baseline (60% ET), 120% ET, and 180% ET, collected during the 2019 season. Values are means ± SE (n=9). P < 0.05; significance was evaluated by one-way analysis of variance and Tukey HSD was used to study post hoc differences among treatments, letters denote significant differences.

**Table S3.** Midday stem water potentials (Ψ _stem_) from vines with three differential irrigation treatments during HWs, Baseline (60% ET), 120% ET, and 180% ET, collected during the 2020 season. Values are means ± SE (n=9). P < 0.05; significance was evaluated by one-way analysis of variance and Tukey HSD was used to study post hoc differences among treatments, letters denote significant differences.

**Table S4.** Midday stem water potentials (Ψ _stem_) from vines with three differential irrigation treatments during HWs, Baseline (60% ET), 90% ET, and 120% ET, collected during the 2021 season. Values are means ± SE (n=9). P < 0.05; significance was evaluated by one-way analysis of variance and Tukey HSD was used to study post hoc differences among treatments, letters denote significant differences.

**Table S5.** Midday measurements of IRT leaf temperatures from vines with three differential irrigation treatments during HWs, Baseline (60% ET), 120% ET, and 180% ET, carried out throughout the 2019 season. Values are means ± SE (n=9). P < 0.05; significance was evaluated by one-way analysis of variance and Tukey HSD was used to study post hoc differences among treatments, letters denote significant differences.

**Table S6.** Diurnal measurements of leaf net photosynthesis (A) carried out during HW2_2019_ on August 15^th^, 2019 of vines exposed to three differential irrigation treatments during HWs, Baseline (60% ET), 120% ET, and 180% ET. Values are means ± SE (n=9 for predawn measurements and for the rest of diurnal measurements n=27). P < 0.05; significance was evaluated by one-way analysis of variance and Tukey HSD was used to study post hoc differences among treatments, letters denote significant differences.

**Table S7.** Diurnal measurements of leaf stomatal conductance (g_s_) carried out during HW2_2019_ on August 15^th^, 2019 of vines exposed to three differential irrigation treatments during HWs, Baseline (60% ET), 120% ET, and 180% ET. Values are means ± SE (n=9 for predawn measurements and for the rest of diurnal measurements n=27). P < 0.05; significance was evaluated by one-way analysis of variance and Tukey HSD was used to study post hoc differences among treatments, letters denote significant differences.

**Table S8.** Diurnal measurements of leaf net photosynthesis (A) carried out during HW2_2020_ on July 12^th^, 2020 of vines exposed to three differential irrigation treatments during HWs, Baseline (60% ET), 120% ET, and 180% ET. Values are means ± SE (n=9 for predawn measurements and for the rest of diurnal measurements n=27). P < 0.05; significance was evaluated by one-way analysis of variance and Tukey HSD was used to study post hoc differences among treatments, letters denote significant differences.

**Table S9.** Diurnal measurements of leaf stomatal conductance (g_s_) carried out during HW2_2020_ on July 12^th^, 2020 of vines exposed to three differential irrigation treatments during HWs, Baseline (60% ET), 120% ET, and 180% ET. Values are means ± SE (n=9 for predawn measurements and for the rest of diurnal measurements n=27). P < 0.05; significance was evaluated by one-way analysis of variance and Tukey HSD was used to study post hoc differences among treatments, letters denote significant differences.

**Table S10.** Diurnal measurements of IRT leaf temperature carried out during HW2_2019_ on August 15^th^, 2019 of vines exposed to three differential irrigation treatments during HWs, Baseline (60% ET), 120% ET, and 180% ET. Values are means ± SE (n=9). P < 0.05; significance was evaluated by one-way analysis of variance and Tukey HSD was used to study post hoc differences among treatments, letters denote significant differences.

**Table S11.** Diurnal measurements of IRT leaf temperature carried out during HW2_2020_ on July 12^th^, 2020 of vines exposed to three differential irrigation treatments during HWs, Baseline (60% ET), 120% ET, and 180% ET. Values are means ± SE (n=9). P < 0.05; significance was evaluated by one-way analysis of variance and Tukey HSD was used to study post hoc differences among treatments, letters denote significant differences.

**Table S12.** Diurnal measurements of leaf transpiration rate (E) carried out during HW2_2019_ on August 15^th^, 2019 of vines exposed to three differential irrigation treatments during HWs, Baseline (60% ET), 120% ET, and 180% ET. Values are means ± SE (n=9 for predawn measurements and for the rest of diurnal measurements n=27). P < 0.05; significance was evaluated by one-way analysis of variance and Tukey HSD was used to study post hoc differences among treatments, letters denote significant differences.

**Table S13.** Diurnal measurements of leaf transpiration rate (E) carried out during HW2_2020_ on July 12^th^, 2020 of vines exposed to three differential irrigation treatments during HWs, Baseline (60% ET), 120% ET, and 180% ET. Values are means ± SE (n=9 for predawn measurements and for the rest of diurnal measurements n=27). P < 0.05; significance was evaluated by one-way analysis of variance and Tukey HSD was used to study post hoc differences among treatments, letters denote significant differences.

**Table S14.** Time series of daily maximum and minimum soil temperature for each of the treatments, baseline (60% ET), 120% ET, and 180% ET, at a depth of 5 cm during the 2020 season. Soil temperature readings for the replicate blocks (n=3) were averaged and confidence intervals calculated from replicates.

**Table S15.** Time series of daily maximum and minimum soil temperature for each of the treatments, baseline (60% ET), 90% ET, and 120% ET, at a depth of 5 cm during the 2021 season. Soil temperature readings for the replicate blocks (n=3) were averaged and confidence intervals calculated from replicates.

**Table S16.** Time series of daily mean and maximum VPD for each of the treatments, baseline (60% ET), 120% ET, and 180% ET, during the 2020 season. VPD was calculated using air temperature and relative humidity sensors located within the vine canopy at one station per treatment.

**Table S17.** Time series of daily mean and maximum VPD for each of the treatments, baseline (60% ET), 180% ET, and 120% ET, during the 2021 season. VPD was calculated using air temperature and relative humidity sensors located within the vine canopy at one station per treatment.

**Table S18.** Dynamics of berry primary metabolites during ripening in the 2019 season of treatments exposed to differential irrigation (Baseline (60% ET), 120% ET, and 180% ET). Total soluble solids expressed as Brix, pH, and titratable acidity expressed in g/L of tartaric acid equivalents. Values are means ± SE (n=9).

**Table S19.** Dynamics of berry primary metabolites from fruitset to harvest in the 2020 season of treatments exposed to differential irrigation (Baseline (60% ET), 120% ET, and 180% ET). Total soluble solids expressed as Brix, pH, and titratable acidity expressed in g/L of tartaric acid equivalents. Values are means ± SE (n=9).

**Table S20.** Dynamics of berry primary metabolites from fruitset to harvest in the 2021 season of treatments exposed to differential irrigation (Baseline (60% ET), 90% ET, and 120% ET). Total soluble solids expressed as Brix, pH, and titratable acidity expressed in g/L of tartaric acid equivalents. Values are means ± SE (n=9).
