## Supplementary material for "Optimizing irrigation during heat events sustains grapevine physiology and fruit production": Survey Results Summary

**Heatwave Responses Survey results summary: 2020**

- Obtained IRB approval (i.e., exemption) on March 13, 2020 via email
- Sent Google Forms survey in April 2020 and collected responses through September 2020
  - Majority of responses came in June 2020
- 58 responses
- 34.5% have their primary operations in Southern California; 24.1% are primarily in the Central Valley; 24.1% are primarily in the Sierra Foothills; and 10.3% are primarily in the North Coast region.
- The majority of respondents (69.6%) own, manage, or oversee fewer than 50 acres, while nearly 20% (19.6%) own, manage, or oversee 500 acres or more.
- The majority (58.6%) of respondents are solely responsible for making management decisions in the vineyard
- The most commonly identified *primary* grape varieties grown by respondents are cabernet sauvignon (37.9%), Chardonnay (12.1%), and Petit Syrah (8.6%).
  - Note that respondents were only allowed to choose *one* grape variety
- Of the climate/weather issues that were most reported as a “moderate” “serious” or “devastating” problem by at least half of the respondents:
  - Drought = 46 respondents
  - Increased pest / disease pressure = 36 respondents
  - Extreme daytime temperatures = 45 respondents
    - This was also reported by the greatest number of respondents (n=8) as being a “devastating” problem (the “highest” or “worst” in the rank), followed by drought (n=7)
  - Warmer nighttime temperatures = 31 respondents
- Only 14.5% of respondents believe that increased water application during the most extreme heat waves in their memory was highly effective at minimizing crop damages
- More than half (54.5%) believe that increased water application was moderately effective at minimizing crop damages, and nearly a quarter (23.6%) felt that increased water application was only slightly effective.
- 8.6% of respondents do not feel prepared to mitigate damages from heatwaves / extreme heat events. Only 10.3% feel extremely prepared. The majority (53.4%) feel moderately prepared, and a little more than a quarter (27.6%) feel slightly prepared.

Scenario for following question responses: *You are informed through your most trusted weather forecasting source that a 3-day heatwave will be impacting your principal growing region identified in Question 1. Assuming nighttime temperatures fall by at least 30°F from daily highs, at what forecasted high temperature would you choose to increase irrigation to your primary grape variety in an effort to mitigate damages from the heatwave?*

- When asked above what forecast temperature would you increase irrigation during veraison, 41.4% of respondents indicated that they would increase irrigation at forecast temperatures above 100F, while 22.4% and 17.2% would increase irrigation above forecast temperatures of 95F and 90F, respectively. Meaning that at temperatures of 100F, more than 80% of respondents have elected to increase their irrigation.
  - Interestingly, half of the respondents indicated that if the temperature was forecast to be *above* the threshold at which they’d elect to increase irrigation they’d only make a small (<30%) additional increase in irrigation. Though 41.4% said they would make a moderate (30-70%) increase
- More than three quarters (76.8%) indicated that they would increase their irrigation before the event. 37.5% of respondents indicated that they would also increase their irrigation during the heat event.
- The temperature thresholds at which growers would elect to increase their irrigation during harvest were more varied. 17.5% would *not* increase irrigation. 19.3% have a threshold of 90F, 19.3% have a threshold of 95F, and 29.8% have a threshold of 100F
  - As with heat events during veraision, most (54.4%) said they would only increase their irrigation by a small (<30%) amount if temperatures were forecast to exceed their threshold. Unlike with veraison, where about 95% of respondents indicated they would increase their irrigation to some degree, during harvest 22.8% indicated they would not increase their irrigation at all
    - Note that 22.8% “no increase” does not align with the 17.5% “no increase” response from the prior question – I suspect this is coming from the people who responded to the “what’s your threshold temperature” question with some variation on “it depends”
- As for timing, heat events during harvest still show folks increasing their irrigation primarily before the event (66.1%) and during the even (32.1%)
  - Note that for this question (and the timing question above) respondents could choose all that apply.
