## Supplementary figures and images for "Optimizing irrigation during heat events sustains grapevine physiology and fruit production"

### Figure S1

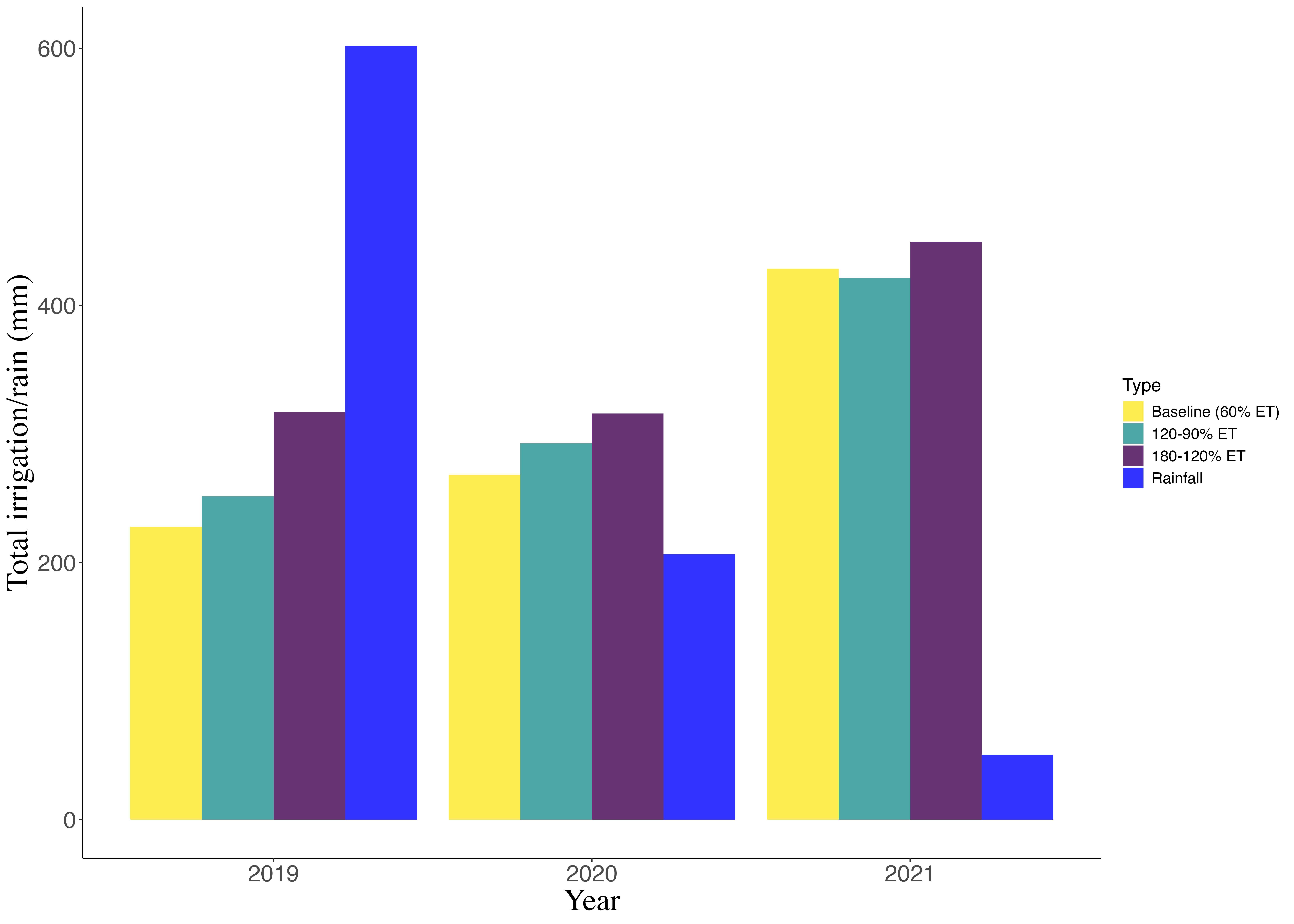
